# Quantifying the success of prey crypsis, aposematism and evasiveness in avoiding predators’ attack

**DOI:** 10.1101/2024.10.31.621260

**Authors:** Daniel Linke, Jacqueline Hernandez Mejia, Valery N. P. Eche Navarro, Prapti Gohil, César Ramírez García, Letty Salinas, Marianne Elias, Pável Matos-Maraví

## Abstract

Antipredator defences typically act at distinct stages of the predation sequence—encounter, identification, approach, and subjugation. However, their effectiveness has rarely been quantified and compared simultaneously in wild predator-prey systems. We conducted a study in Peru, where we installed aviaries at two localities and recorded the responses of wild avian predators to three types of antipredator defences—crypsis, aposematism, and evasiveness—expressed by three butterfly species. The study included both immature and adult birds from insectivorous species in forest and urban environments. We tested the theoretical expectations that cryptic butterflies (Nymphalidae: Euptychiina) were rarely detected, aposematic *Heliconius* (Nymphalidae: Heliconiinae) were often sighted but seldom attacked, and evasive *Spicauda* (Hesperiidae: Eudaminae) were frequently detected and attacked but evaded capture at higher rates. Despite these distinct defensive strategies, mortality rates among prey types were largely similar, but predator life stage strongly influenced defence effectiveness. Immature birds were more likely to attack *Heliconius*, possibly reflecting a lack of learned avoidance for aposematic signals. Additionally, predator family influenced predation patterns, with more skilled insectivores (e.g., Vireonidae) showing higher capture success against defended prey. These findings illuminate the evolutionary pressures that shape predator behaviour and prey defences in tropical ecosystems, where high predator diversity possibly maintains the coexistence of multiple defensive strategies. The observed similar mortality rates underscore the adaptive value of these defences, as they collectively distribute the total predation pressure across prey species.

## Introduction

Predation selective pressure have shaped the evolution of a wide range of defensive strategies in prey species. Antipredator mechanisms are often best fit at deterring one of the sequential stages of the predation sequence (Endler, 1991; Ruxton et al., 2018). For example, crypsis reduces the likelihood of encountering and detection, whereas aposematism relies on conspicuous signals that advertise unprofitability to deter attacks by predators that learned the warning signals. In contrast, evasive ability is a defence strategy that impede subjugation and consumption once predators engaged in prey pursuing. However, understanding how wild predators interact with and respond to the occurrence of multiple defended prey, and how the effectiveness of antipredator defences is shaped by the community composition of predators, provide crucial insights into the evolutionary pressures affecting prey phenotypic diversity.

Among prey species exhibiting a broad arsenal of defensive strategies, butterflies possess antipredator defences that operate on each stage of the predation sequence. Cryptic butterflies, for example, typically reduce predator encounters and detection through background matching (Feltmate & Williams, 1989; Vallin et al., 2006; Pinheiro & Campos, 2019; Pembury Smith & Ruxton, 2021), which can be assessed in the wild by wing damage, though the true predation risk of cryptic prey remains unclear (Molleman et al., 2020). Unpalatable butterflies, such as Heliconiini and Ithomiini (Mallet & Gilbert, 1995), advertise their unprofitability through aposematic cues that avian predators learn to avoid (Arias et al., 2016; Chouteau et al., 2016; Willmott et al., 2017; Chouteau et al., 2019), but experiments struggle to assess rejection after an initial attack (Mappes et al., 2014; Seymoure et al., 2018). Finally, evasive flight behaviour (Chai, 1986; Srygley & Chai, 1990; Pinheiro & Freitas, 2014) and deflective structures may prevent deadly attacks to butterflies after being spotted (Chotard et al., 2022). Evasiveness triggering predator learning and being used during prey identification is theoretically possible (Ruxton et al., 2004), and recent studies have shown predators prefer attacking non-evasive butterfly models (Páez et al., 2021; Linke et al., 2022).

No antipredator defence offers complete protection against predators, especially in ecosystems harbouring a large diversity of predator species. However, our understanding on how wild predators respond to simultaneously encountered prey with diverse defence strategies is limited (Pinheiro & Campos, 2019; wild predators but dummy prey - Guerra et al., 2024; Dell’aglio et al., 2016). By conducting behavioural experiments in aviaries with wild insectivorous birds, we aim to understand how predators respond when preys with distinct antipredator defences are presented simultaneously, as well as how effective different anti-predator defences are when avoiding predation.

The prey that we study are butterflies with antipredator defences believed to act at each major stage of the predation sequence: 1) Euptychiina (Nymphalidae: Satyrinae), which likely relies on crypsis to avoid being detected, 2) *Heliconius* (Nymphalidae: Heliconiinae), which likely relies on aposematism and Müllerian mimicry to avoid being identified as profitable prey, and 3) *Spicauda* (Hesperiidae: Eudaminae), which likely relies on evasive behaviour to avoid being caught. We investigate how predator responses vary by life stage, diet, and taxonomic family.

We postulate the following predictions stemming from predator-prey-interaction theory:

**(I) Probability of remaining undetected:** cryptic butterflies (i.e., the dull brown colouration of Euptychiina and *Spicauda*) have lower probabilities to be detected by birds compared to a conspicuous butterfly (i.e., the aposematic *Heliconius*).
**(II) Probability of being rejected upon identification:** butterflies that advertise unprofitability by the means of conspicuous colour patterns (i.e., *Heliconius*) have a higher probability of being sight rejected, without active attacking behaviour, by predators that have learned the visual cues associated with unprofitable prey (as opposed to, e.g., the presumably slow flying, dull coloured and palatable Euptychiina).
**(III) Probability of capture avoidance upon predator attack:** butterflies with evasive flight behaviours (i.e. *Spicauda*) are more likely to escape predator attacks compared to slow-flying butterflies (e.g., Euptychiina or *Heliconius*).
**(IV) Probability to survive upon predator attack:** butterflies with structural/chemical defences (such as the sturdy wings of unpalatable *Heliconius* or the hindwing tails of *Spicauda*) survive predator attacks by being released with non-fatal damage.

## Methods

### Study location

The experiments were carried out during the rainy season of 2021/22 (October – February) and the dry season of 2022 (June – September). The two study locations were 1) within the city limits of Tarapoto, Peru (6.47°S, 76.35W, ∼400 masl), and 2) within a privately protected area, the Urahuasha sector (6°27′S 76°20′W, ∼700 masl), near the external border of the “Área de Conservación Regional Cordillera Escalera”, San Martin, Peru. The first study site was characterised by significant human intervention, and the second site was characterised by secondary growth forest (∼50 years old) with very limited human influence.

### Birds

We installed dull-coloured mist nets near pathways in scrubs, secondary forests, forest edges and hill crests at both study sites (Supplement 1). Nets were opened since ∼7 AM and checked hourly until dusk (∼5 PM). Captured birds were identified to species, their age determined (immature or adult) and photographed. Immature birds were identified by plumage and commissure colour. Birds selected for behavioural experiments were known to feed on butterflies (Pinheiro & Cintra, 2017), classified as insectivorous (Egg et al., 2010) or their insectivorous diet confirmed by local experts. Birds were categorized as fully insectivorous or omnivorous/insectivorous. All other birds were marked and released. Experimental birds were kept in captivity for up to 30 hours at the urban site and 12 hours at the forest site, with constant access to water and local papaya.

### Butterflies

**(I)** As representatives of cryptic prey, we studied medium-sized Euptychiina, consisting of mainly *Cissia penelope* and related, syntopic species in the genera *Cissia*, *Magneuptychia* and *Yphthimoides.* **(II)** As representatives of unpalatable, aposematic prey, we studied *Heliconius erato*, and when unavailable, we used the co-mimetic *H. melpomene*. **(III)** As representatives of prey relying on evasive flight to avoid predation, we studied *Spicauda simplicius*, and when unavailable, we used the morphologically similar and syntopic *S. teleus*, *S. tanna*, and *S. procne*. All species within each study phenotype shared similar morphologies, behaviours, and co-occurred in the same microhabitats.

Butterflies were caught with entomological nets around the study sites and identified to the species level, with Euptychiines sometimes only to the genus level. They were kept alive in mesh cages (50×50×50 cm) with fake flowers filled with sugar water. Each butterfly was used in up to three predation experiments or for a maximum of 48 hours to prevent fatigue. Afterward, they were sacrificed and stored in silica gel for morphological identification.

### Behavioural experiments in aviaries

At each study location and in open space to obtain similar luminosity, we constructed an experimental aviary (H×W×L = 4×2×4 m) made of green garden mesh and plumbing tubes (Supplement 2). We installed bamboo perches for the birds to rest on at ∼50 cm below the aviary ceiling. We installed two video cameras (GoPro HERO6, San Mateo, USA; set to 1080p, 60 fps and the wide lens setting) on the two front sides of the aviary at ∼1 m height, allowing for the complete video capture of the inside of the aviary. The experiments were conducted from ∼7 AM to ∼6 PM depending on availability of birds and butterflies.

The behavioural experiments involved simultaneously presenting the three butterfly types to a bird in the aviary and recording the predator and prey interactions. After starving for 60 minutes, the bird was acclimatized in the aviary for 30 minutes. Then, the butterflies were released and the cameras recorded the experiment for 60 minutes, while avoiding any human presence to reduce stress. At the end, any remaining butterflies were recaptured and the ground was surveyed for any detached wings. After cutting two tail feathers for identification, the bird was released before dark in favourable weather. Recaught birds were not used again. The experiment was invalid if there were disturbances like sudden rainfall, the appearance of cats or monkeys, or signs of bird fatigue (e.g., lack of movement or erratic flight).

### Video processing

We aligned the two cameras to allow for the simultaneous observation of both cameras, using DaVinci Resolve 18 (Blackmagic Design, Melbourne, Australia). We noted the behavioural responses of birds and butterflies as timestamps (Figure 1), recording the start and end of stages in the predation sequence: encounter, identification, approach, subjugation and consumption. Each bird-butterfly combination was assigned the highest observed stage in the predation sequence. Birds that did not interact with any butterfly were excluded from analysis (i.e., no prey sight-rejection or attack). We also recorded the approximate time the bird and each butterfly co-occurred at two heights in the aviary: ground level, 0–200 cm, and ceiling level, 200–400 cm.

**Figure 1:**
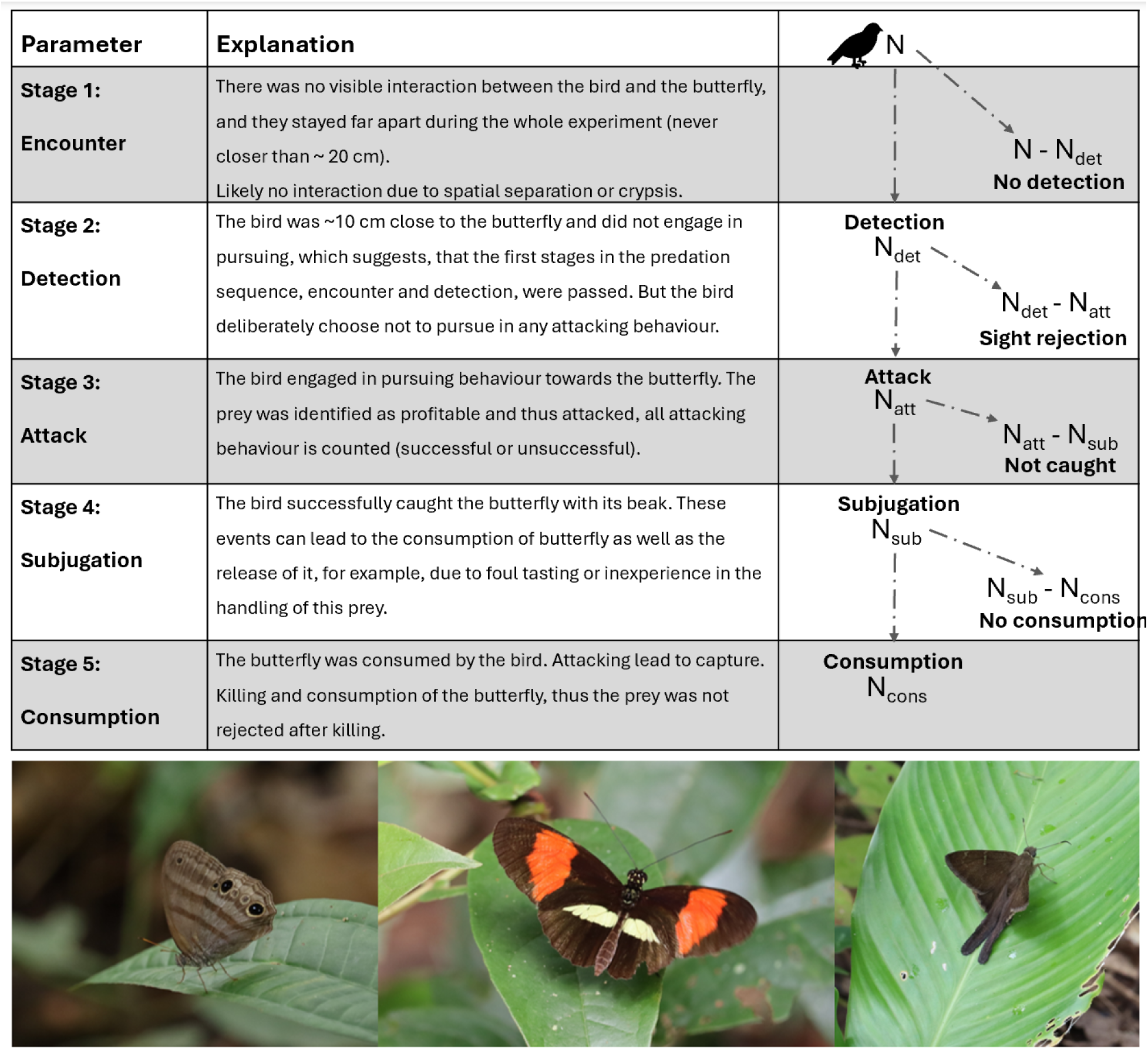
(above) Stages of the predator-prey-interactions observed during the experiment, including a dichotomous flowchart depicting the progression of birds through the predation sequence. (below) Examples of butterflies in their native environment, used in the aviary experiments to assess responses of birds to different anti-predation strategies, from left to right: the cryptic Cissia penelope (Fabricius, 1775), the aposematic Heliconius erato (Linnaeus, 1758) and the evasive Spicauda simplicius (Stoll, 1790) (photos not scaled).

### Chance of predator-prey encounter

To estimate the encounter probability, we calculated the co-occurrence rate of the bird and each butterfly, using:

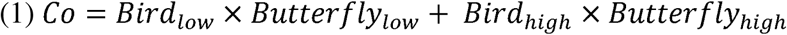

Where, the first term is the proportion of experimental time the bird and a butterfly co-occurred in the ground level (0– 200 cm) and the second term is the proportion of time they co-occurred in the ceiling level (200–400 cm). The *Co* index, ranges between 0 and 1, where 0 is a complete spatial mismatch between the bird and a butterfly (very unlikely the predator encounters the prey) and 1 is a complete overlap (highly likely that the predator encounters the prey).

### Predictors of bird responses to prey phenotypes: logistic regressions

We used generalized linear mixed models (GLMMs) with a binomial distribution (via the *glmer* function from the lme4 R package; Bates et al., 2015) to identify bird responses (sight rejection or attacking) in relation to prey type, bird age (immature or adult), and diet (insectivorous or mixed). Bird identity was included as a random effect to account for individual variability. Each bird-butterfly interaction was transformed into a binary outcome based on the highest observed behavioural stage (Figure 1). These two separate models were used to compare sight rejection and attack behaviours:

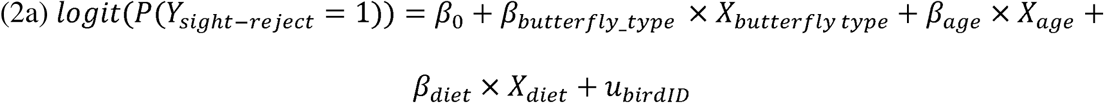

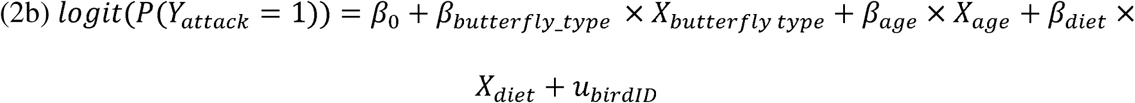

Where Y is the binary outcome of a butterfly to be sight rejected or attacked by a bird;, B_0_ is the general tendency to sight reject or attack;, *β_butterfly_type_* x *X_butterfly type_*, *β_age_* x *X_age_* and, *β_diet_* x *X_diet_* represents the effects of butterfly type, age and diet, respectively, and *u_bird!D_* accounts for variation among individual birds. For the sight-rejection model (2a), the response variable was coded as 1 for sight rejection (N_det_ – N_att_, i.e., by being ∼10 cm close to the butterfly without attacking behaviour) and 0 for attacking behaviour (N_att_). In the attacking model (2b), the response variable was coded as 1 for active attacking (N_att_), and 0 for all other responses (N_det_ – N_att_ and N – N_det_, i.e., sigh rejection or no interaction at all). Additional models incorporating season, locality, and bird family are described in Supplement 3.

### Probabilities of attack deterrence at different stages in the predation sequence

We used log-likelihood tests to compare the probabilities of a prey to (1) remain undetected, (2) get sight-rejected upon detection, (3.1) get attacked, (3.2) avoid capture during a predator attack and (4) survive a predator attack. The tests were performed using the birds that detected at least one butterfly prey (any interaction: N – N_det_), i.e., birds that were ∼10 cm close to the prey and birds that showed active attacking behaviour (Figure 1). Further tests were also performed using the data subset by predator age (immature or adult) (additional models for season, wet or dry, and for habitat, urban or forest, in Supplement 4). We calculated the log-likelihood of models depicting the number of events within each predation sequence stage for a particular prey phenotype compared to others, following the method implemented in Mérot et al. (2015), Willmott et al. (2017) and Páez et al. (2021):

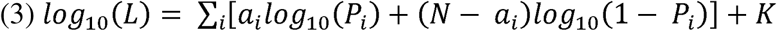

Where *i* represents the prey type (Euptychiina, *Heliconius* or *Spicauda*); *N* denotes the total count the prey was presented, *a_i_* is the count of interactions with the prey type *i*; *P_i_* is the attack probability for each prey type *i*; and *K* is a constant that cancels out when comparing models. For each stage, we compared five competing models: all prey having different probabilities of interaction (“all different”), the same probabilities (“all equal”), and two of the prey phenotypes having the same probability but different from the third phenotype (“Euptychiina different”, “*Heliconius* different” and “*Spicauda* different”). The model with the lowest AICc (corrected Akaike information criterion) value was selected as the best model, while any model within 2 units of AICc was not rejected. We calculated Akaike weights following Burnham & Anderson (2013) to compare differences between the five competing models for each predation stage.

### Description of the likelihood-based model tests for each stage in the predation sequence

**(1) Probability to remain undetected:** We compared the sum of occasions for each prey phenotype where the bird and butterfly did not interact (N – N_det_, i.e., prey was not encountered or detected) to the total number of tested birds (N).

In addition to the likelihood-based comparison, we compared *Co* index values between butterflies using a pairwise Wilcoxon signed-rank tests with Bonferroni correction (for multiple comparisons) as normal distribution of the data was rejected (Shapiro-Wilk test: N = 393, W = 0.897, p = 1.355e^−15^).

**(2) Probability of being rejected upon detection:** We compared the sum of events for each prey phenotype where the bird was close to the butterfly (∼10 cm) but did not engage in any attacking behaviour (N_det_ – N_att_), in comparison to all birds that did enter the predation sequence for at least one butterfly (N). We assumed that the birds in spatial proximity to a butterfly were able to spot the prey but decided not to pursue any attacking behaviour (sight rejection).

**(3) Targeting probability and capture avoidance:** This involves two steps, **(3.1)** the chance of getting attacked in the first place and **(3.2)** the ability to avoid being caught when attacked. First, **(3.1)** we compared the sum of events where the butterfly was actively attacked by the bird depending on the butterfly type, regardless of the outcome (N_att_), in comparison to all birds that entered the predation sequence with this prey phenotype (N_det_). Second, **(3.2),** to determine whether prey phenotypes differ in their ability to avoid being captured, we additionally computed the sum of events where the butterfly was not captured (N_att_ – N_sub_) in comparison to the sum of birds showing active attacking behaviour to each prey phenotype (N_att_).

Lastly, we determined for each bird and butterfly combination the time for the first instance of attacking behaviour (no attack, equals 60 minutes) to compare any preference in engaging to pursue different butterfly types. As a normal distribution of the data was rejected (Shapiro-Wilk test: N = 217, W = 0.747, p = 2.2e^−16^), we performed pairwise Wilcoxon signed-rank tests with Bonferroni correction (for multiple comparisons) to compare initial attack time distributions between butterflies.

**(4) Probability to survive upon predator attack:** We compared the sum of failing to kill (N_att_ – N_cons_) for each prey phenotype in comparison to the number of actively attacked butterflies per phenotype (N_att_).

Statistics (except log-likelihood tests, which were implemented in a spreadsheet; Supplement 4) were done using R v. 4.3.0 (R Core Team, 2023). Graphics were generated using the R package *ggplot2* v. 3.4.3 (Wickham, 2016).

## Results

We performed 262 experiments, of which 216 were deemed valid. There were 65 valid experiments in the urban environment and 151 experiments in the forested habitat, of which 70 were performed in the wet season and 81 in the dry season. Due to time and logistics constraints, we were not able to carry out experiments in the urban environment during the dry season.

We studied 41 bird species from 13 families. The number of insectivorous species was higher in the forested habitat than in the urban environment (urban/wet season = 12 species; forest/wet season = 19 species; forest/dry season = 23 species) (Supplement 5). Complete bird lists and photographs are available in Field Museum guides (Navarro et al., 2023, Hernández Mejía et al. *in review*). In the urban area, 68% of birds were adults and 32% were immatures, while in the forest, 29% were adults and 71% immatures during the wet season, and 16% adults and 84% immatures in the dry season (species-level data in Supplement 5).

### General differences between prey types and bird families

Of birds that sight rejected or attacked at least one butterfly, nearly 70% did not interact with Euptychiines (N – N_det_), whereas this proportion was notably lower for *Heliconius* (34%) and *Spicauda* (18%). Sight rejection (N_det_ – N_att_) was most prevalent toward *Heliconius*, with almost 50% of birds exhibiting this response, followed by *Spicauda* (18%) and Euptychiines (9%). *Spicauda* was most frequently attacked by birds (N_att_, 40%), while attack rates for Euptychiines and *Heliconius* were comparable at 21% and 18%, respectively. Only 6% of all birds caught *Heliconius* (i.e., N_sub_, killing or releasing it afterwards), compared to 16% for *Spicauda* and 12% for Euptychiines (Supplement 6). Overall mortality rates (N_cons_) were similar across all three prey phenotypes, with Euptychiines at 7%, *Heliconius* at 4%, and *Spicauda* at 9%.

Among bird families, Furnariidae, were most effective in attacking Euptychiines, with 21% successfully catching them, while Thraupidae were the least effective, with only 8% exhibiting some interaction (i.e., at least sight rejection). Bird families showed similar probabilities of sight rejecting *Heliconius*, ranging between 30-50%. Tyrannidae and Vireonidae, two highly insectivorous families and skilled flycatcher hunters, were the only birds that did not sight reject *Spicauda*, with Vireonidae displaying the highest success in capturing them (Supplement 6).

### Predictors of bird behaviours to prey phenotypes: logistic regressions

The GLMM for sight rejection behaviour revealed that birds more frequently rejected *Heliconius* than Euptychiines (z = 7.04, p = 1.94e^−12^), while there were no significant differences for *Spicauda* (z = −0.15, p = 0.882), bird age (z = 0.71, p = 0.479), or diet (z = 0.33, p = 0.745). There was substantial variability across individuals revealed by the random intercept variance (σ^2^ = 305.2, SD = 17.47). In the GLMM for attacking behaviour, birds were significantly more likely to attack *Spicauda* than Euptychiines (z = 3.56, p = 0.0008). No significant differences were observed for *Heliconius* (z = −0.83, p = 0.408), bird age (z = −0.50, p = 0.617), or diet (z = 0.39, p = 0.699). Individual variation in attacking behaviour was low (σ^2^ = 0.88, SD = 0.94). Additional GLMM models including bird family, habitat and season can be found in Supplement 3. Notably, in such GLMM models, Vireonidae birds were significantly less likely to sight-reject (z = −2.19, p = 0.028) and more likely to attack prey butterflies (z = 2.11, p = 0.035) compared to other bird families. This interaction was barely significant for Tyrannidae (sight rejection, z = −1.67, p = 0.095, attacking behaviour, z = 1.87, p = 0.062).

### Likelihood-based comparisons along the predator-prey interactions

Results for log-likelihood tests for the total dataset, immatures and adults can be found in Figure 2 and Table 1. Tests for habitat (forest vs. urban) and season (dry vs. wet) can be found in Supplement 4. Graphs for season and habitat can be found in Supplement 7.

**Figure 2:**
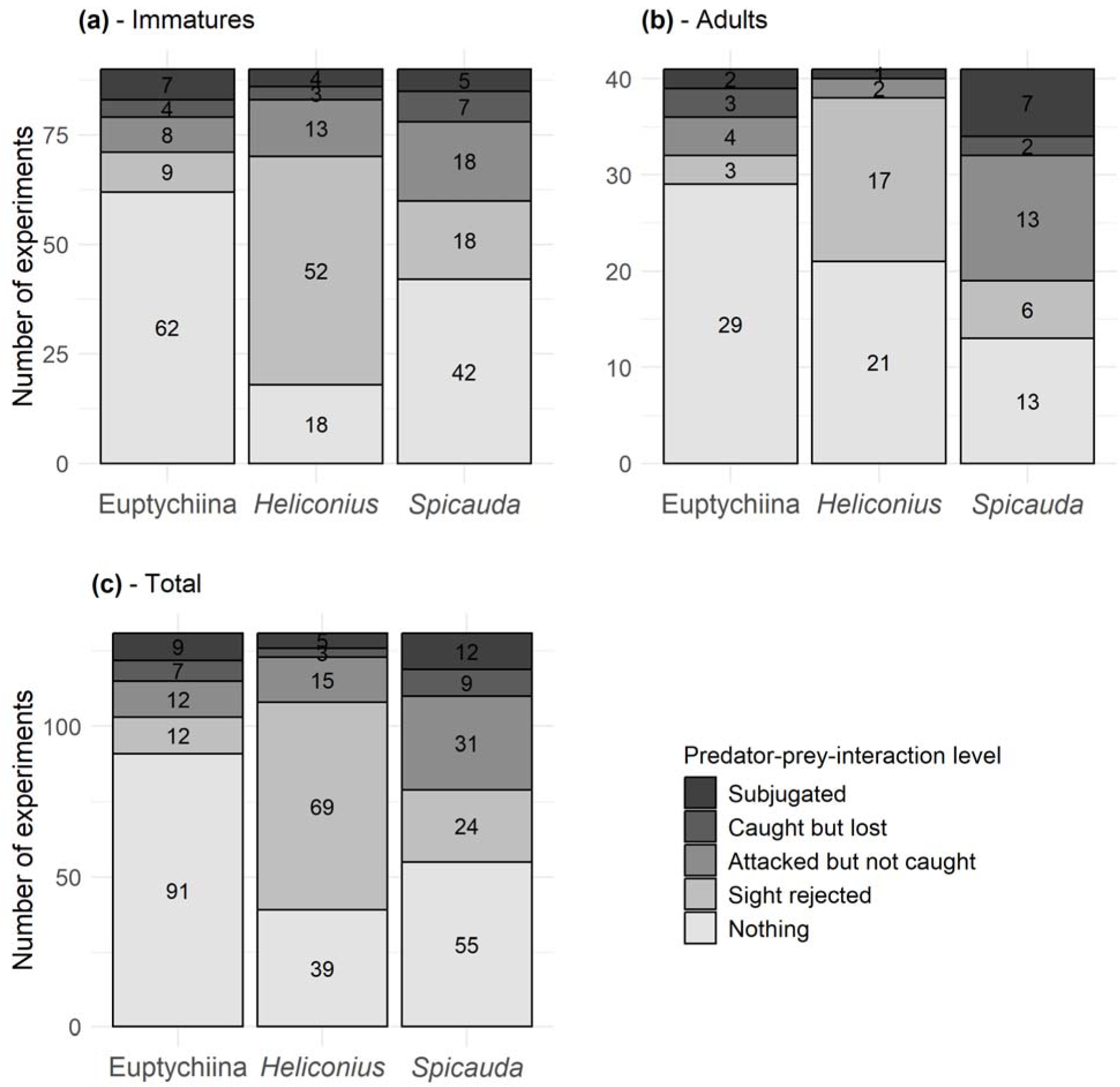
Comparison of predation outcomes (no interaction, sight rejected, attacked, caught or subjugated) depending on the butterfly type (Euptychiines, Heliconius and Spicauda) and depending on the age of experimental birds (adult and immature).; **(a)** distribution for all immature birds, **(b)** distribution for all adult birds and **(c)** distribution of all predator-prey interactions including all valid experiments.

**Table 1:** Likelihood-based model tests of scenarios depicting the effectiveness of antipredator defences by three butterfly prey along the predation sequence. Akaike weights for the total number of birds and bird age (immature vs. adult) are shown. Best models are marked in **<u>bold and underlined</u>**, while competing models (ΔAICc < 2) are only **bold**. Values in brackets represent ΔAICc values compared to the best performing model. Additional results for seasons and habitats can be found in Supplement 4.

| Stage | Model | All different | All equal | *Heliconius*<br>different | *Spicauda*<br>different | *Euptychiina*<br>different |
| --- | --- | --- | --- | --- | --- | --- |
| (1) Remaining undetected | Total | **0.47** (0.18) | 0.00 (15.45) | 0.02 (6.98) | 0.00 (16.58) | **<u>0.51</u>** (0.00) |
|  | Immature | **<u>0.67</u>** (0.00) | 0.00 (15.80) | **0.25** (1.95) | 0.00 (17.77) | 0.08 (4.34) |
|  | Adult | **0.22** (0.69) | **0.11** (2.10) | 0.04 (4.17) | **0.31** (0.03) | **<u>0.32</u>** (0.00) |
| (2) Rejection upon identification | Total | **<u>0.50</u>** (0.00) | 0.00 (26.34) | **0.50** (0.02) | 0.00 (25.15) | 0.00 (13.10) |
|  | Immature | **0.44** (0.49) | 0.00 (20.40) | **<u>0.56</u>** (0.00) | 0.00 (19.90) | 0.00 (10.57) |
|  | Adult | **0.24** (1.61) | 0.06 (4.23) | **<u>0.53</u>** (0.00) | 0.03 (5.59) | 0.13 (2.78) |
| (3.1) Targeting probability | Total | **0.25** (1.77) | 0.04 (5.73) | 0.07 (4.29) | **<u>0.62</u>** (0.00) | 0.02 (6.70) |
|  | Immature | 0.10 (2.24) | **<u>0.32</u>** (0.00) | **0.14** (1.68) | **0.29** (0.21) | **0.16** (1.41) |
|  | Adult | **0.36** (0.51) | 0.02 (6.68) | 0.15 (2.31) | **<u>0.47</u>** (0.00) | 0.01 (8.30) |
| (3.2) Capture avoidance | Total | 0.09 (2.89) | **<u>0.37</u>** (0.00) | **0.16** (1.66) | **0.14** (1.88) | **0.24** (0.86) |
|  | Immature | 0.08 (3.30) | **<u>0.39</u>** (0.00) | **0.16** (1.76) | 0.14 (2.01) | **0.22** (1.16) |
|  | Adult | 0.05 (4.37) | **<u>0.46</u>** (0.00) | 0.15 (2.19) | 0.16 (2.15) | **0.17** (1.98) |
| (4) Surviving attack | Total | 0.06 (3.79) | **<u>0.43</u>** (0.00) | **0.16** (1.99) | **0.16** (1.97) | **0.19** (1.67) |
|  | Immature | 0.07 (3.36) | **<u>0.40</u>** (0.00) | **0.15** (2.00) | **0.16** (1.77) | **0.22** (1.18) |
|  | Adult | 0.05 (4.54) | **<u>0.47</u>** (0.00) | 0.15 (2.25) | 0.16 (2.19) | 0.16 (2.12) |

**(1) Probability to remain undetected**: Across all bird families in different habitats and seasons, there was a preference for foraging near the ceiling level (Supplement 8). Antbirds (Thamnophilidae, 23 individuals and 9 species) spent most of the time (66%) foraging near the ground, whereas Vireonidae (9 individuals and 2 species) spent on average only 14% of their time near the ground. Butterflies, on average, differed in their stratum selection (Supplement 8). Euptychiines tended to occur most of the time (55%) near the ground, whereas *Spicauda* and *Heliconius* spent 78% and 83% of their time, respectively, in the ceiling level.

Our likelihood-based model tests supported the “Euptychiines different” scenario (Akaike weight: 0.51; Figure 2, Table 1), which suggested that birds were overall least likely to encounter Euptychiines compared to *Heliconius* and *Spicauda*. Additionally, the “all different” scenario was the second-best model (ΔAICc = 0.18, Akaike weight:0.47), with birds most likely to encounter *Heliconius*. For adult birds, “Euptychiines different” and “*Spicauda* different” had almost equal Akaike weights (0.32 and 0.31 respectively), both scenarios describing that birds were least likely to encounter the Euptychiines and most likely *Spicauda*. However, the “all equal” model (no difference among butterfly species) also had substantial support (Akaike weight: 0.22; ΔAICc = 0.69). For immature birds, “all different” was the best model, with Eptychiines the least and *Heliconius* the most likely to be encountered (Akaike weight: 0.67). Additionally, “*Heliconius* different” received substantial support (ΔAICc = 1.95, Akaike weight: 0.25).

Our non-parametric test of encounter chance for the bird and each butterfly type (*Co* index, Figure 3(a)) suggested that Euptychiines and *Heliconius* (pairwise Wilcoxon signed-rank test, W = 5762; Bonferroni-adjusted p = 1.2e^−5^; Euptychiina’s *Co*: avg. = 0.477, SD = 0.371 and *Heliconius*’ *Co:* avg. = 0.641, SD = 0.344) as well as Euptychiines and *Spicauda* (pairwise Wilcoxon signed-rank test, W = 6648.5; Bonferroni-adjusted p = 0.00071; *Spicauda*’s *Co*: avg = 0.600, SD = 0.351) differed significantly in their spatial co-occurrence, whereas no difference was detected between *Spicauda* and *Heliconius* (pairwise Wilcoxon signed-rank test, W = 9601; Bonferroni-adjusted p = 0.283). This confirmed that while *Heliconius* and *Spicauda* co-occurred in the aviary’s upper part, Euptychiines preferred the ground, reducing their encounter probability.

**Figure 3:**
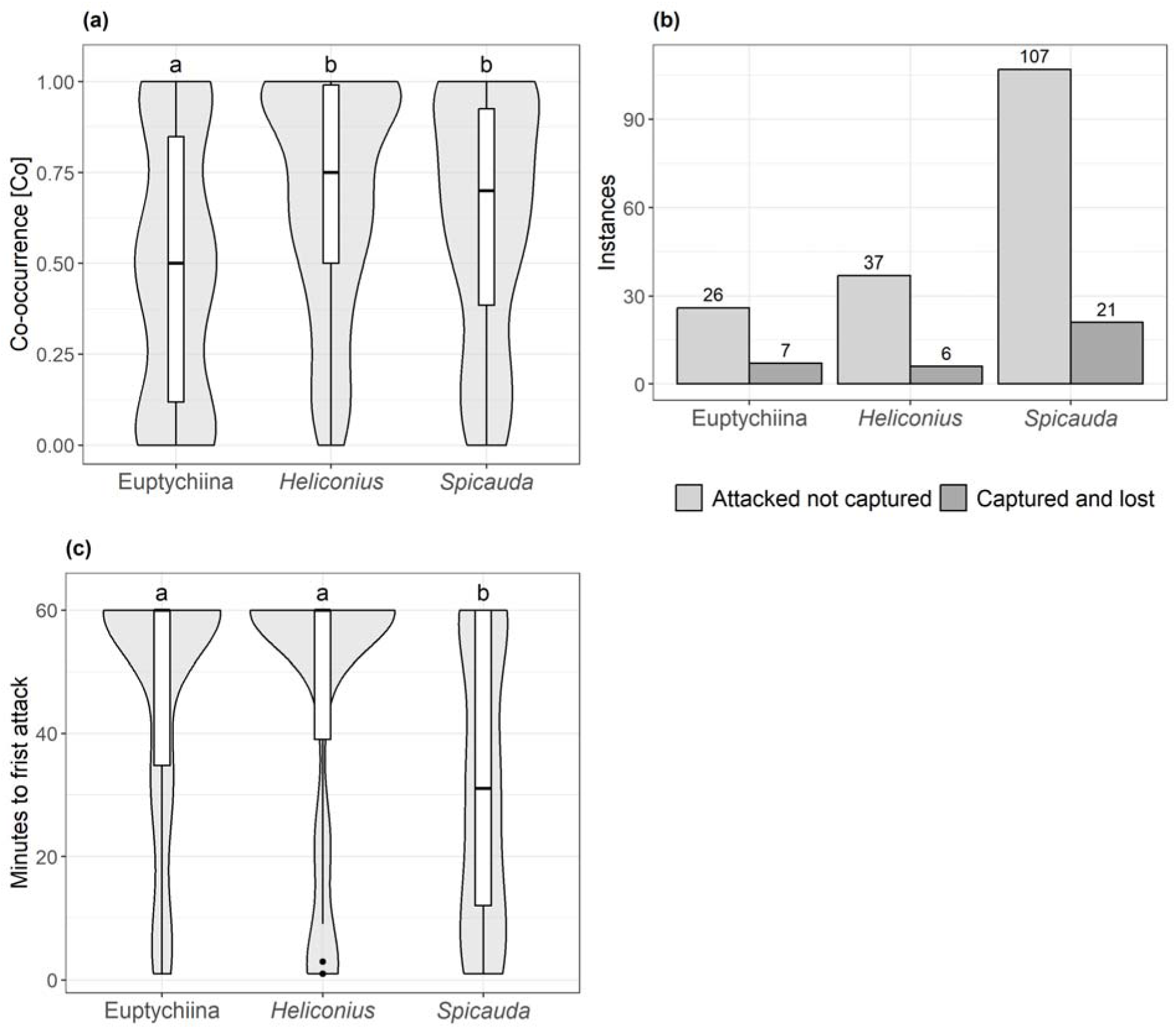
**(a)** Shared occupancy across high and low stratum in the aviary for all three used butterfly types. Compact letters denote significant differences between groups (p<0.05) based on Wilcoxon signed-rank test. **(b)**: Sum of instances of attacking behaviour without capture and capturing but subsequently losing the prey for each butterfly phenotype. **(c)**: Duration until the first attack occurred for this butterfly type within the 60-minute experiment. Only for birds attacking at least one butterfly. Compact letters denote significant differences between butterfly types (Wilcoxon signed-rank test).

**(2) Probability of being rejected upon identification**: The best-selected model was “*Heliconius* different” (Akaike weight = 0.50; Figure 2, Table 1) with “all different” being the second-best model (Akaike weight = 0.50, ΔAICc = 0.02). Both models suggested that most birds sight-rejected *Heliconius*, followed by *Spicauda* and Euptychiines. A similar pattern was supported by both immatures and adults, with immatures exhibiting higher sight-rejection rates towards all butterfly types. *Spicauda,* even though it co-occurred at high rates with most birds in the aviaries, was sight-rejected at a higher rate (the “all different” scenario) than Euptychiines.

**(3.1) Targeting probability:** The “*Spicauda* different” was identified as the best model (Akaike weight = 0.62, Figure 2, Table 1), suggesting that birds more frequently attacked *Spicauda* than Euptychiines and *Heliconius*. This is in line with our observations of birds often attacking *Spicauda* repeatedly until they captured and subjugated it (Figure 3(b)); the ratio of the number of unsuccessful attacks to the sum of all birds attacking a prey is highest for *Spicauda* with 2.05, followed by *Heliconius* with 1.61 and Euptychiines with 0.92. The model “all different” had substantial support (Akaike weight = 0.25, ΔAICc = 1.77), suggesting that Euptychiines were attacked more frequently than *Heliconius* by birds that detected a butterfly prey. For adults, only the “all different” and “*Spicauda* different” models received substantial support (Akaike weights: 0.36 and 0.47, respectively), indicating *Spicauda* as the primary target. In contrast, immatures showed more even attack rates, with “all equal” being the best model (Akaike weight = 0.32) and “all different” was poorly supported (ΔAICc = 2.24, Akaike weight = 0.10), indicating less selective predatorial responses.

**(3.2) Capture avoidance**: “All equal” was the best model, suggesting that all butterflies showed similar likelihood of evading capture upon predator attack (Akaike weight = 0.37, Figure 2, Table 1). For immatures, additional models (“*Heliconius* different,” and “Euptychiines different”) had ΔAICc between 1 and 2 (Akaike weights between 0.16 and 0.22), indicating varying capture efficiency. Conversely, for adult birds only “all equal” and “Euptychiina different” were selected (Akaike weights: 0.46 and 0.17, respectively), pointing to similar capture rates for *Heliconius* and *Spicauda*, with Euptychiines being the easiest to catch To supplement the likelihood-based tests, we compared the times of first predator attack among butterfly types. The pairwise Wilcoxon signed-rank tests suggested that the times to attack differed between *Spicauda* and *Heliconius* (*Spicauda* 32 ± 22.4 min; *Heliconius* 46.6 ± 22.3 min; W = 3627.5; Bonferroni-adjusted p = 9.0e^−05^) as well as between *Spicauda* and Euptychiines (Euptychiina 47.9 ± 19.6 min; W = 3728.5, Bonferroni-adjusted p = 1.4e^−05^) (Figure 3(c)), confirming that *Spicauda* was attacked significantly earlier compared to the other two prey types.

**(4) Probability to survive upon predator attack:** The “all equal” model was preferred (Akaike weight = 0.43), while most other models had ΔAICc > 1.67 and Akaike weights < 0.19. Only the model “all different” was rejected (ΔAICc = 3.79, Akaike weight = 0.06). For immatures, “all equal” was preferred (Akaike weight = 0.40), while the models “*Heliconius* different”, “*Spicauda* different” and “Euptychiina different” were not rejected (ΔAICc: 2.00, 1.77 and 1.18; Akaike weights: 0.15, 0.16 and 0.22, respectively), suggesting that Euptychiines were more likely to be subjugated once a predator attacked it compared to *Spicauda* or *Heliconius*. This pattern was absent for adults, with the “all equal” scenario being the only model with ΔAICc < 2 (Akaike weight: 0.47).

## Discussion

Our study tested four predictions of the predation sequence theory related to the fitness of antipredator defences during the prey encounter, identification, attack and subjugation stages. We studied tropical birds that include insects in their diets and live butterflies displaying antipredator defences (crypsis, aposematism and evasiveness) thought to best function at different stages of the predation sequence (Kikuchi et al., 2023). In controlled aviary settings, we confirm that crypsis reduces the likelihood of encounter and detection, aposematism deters attacks, and evasive flight reduces the probability of subjugation after attacks. However, we also provide evidence that bird age and, to some degree, diet are important predictors of defence effectiveness. Aposematism was less effective against immature birds, while evasive ability was more successful in avoiding attacks from immatures. Finally, we quantify similar survival rates across all predation stages, regardless of antipredator defence. The higher number of insectivorous birds and larger proportion of immatures in the mature secondary forest, compared to the urban environment, were key drivers of the similar mortality rates across butterfly types.

### Defence efficiency along the predation sequence

**Probability to remain undetected:** Cryptic species often remain motionless for prolonged periods of time, are hard to spot (Stevens & Merilaita, 2009; Seymoure et al., 2018; Ruxton et al., 2018) and forage close to the ground (Burd, 1994). Euptychiines are characterized by dull wing colourations and often flying near the ground, which, indeed, resulted in them being the least likely prey phenotype to be encountered by birds in our study. Conversely, while *Spicauda* also has a uniform brown appearance and might be considered cryptic, their conspicuous flight behaviour often renders them in spatial proximity to most avian predators, thus, possibly indicating a relaxed selection for crypsis and a reliance on their evasive flight abilities or hindwing tails to evade predation upon attack (Linke et al., 2024).

We detected a strong preference of fully or partly insectivorous birds towards the upper layer of the aviary (200 - 400 cm). Although the detection rates of prey varied in experiments carried out in the forest and urban environments, this might be due to differences in species specific height preferences of birds or the different abundances of immatures relative to adults in each habitat. Due to limited overlap in species and age distribution between study locations, we cannot determine why prey detection rates shifted across habitats (Supplement 4). Future studies should survey birds over longer periods and in various locations and across habitats, to compare foraging strategies differences among species and life stages. Including other skilled insect-hunting birds in behavioural experiments, such as Jacamars (Galbulidae) (Chai, 1986; Langham, 2006; Pinheiro & Campos, 2019), absent at our study locations, would provide further insights on antipredator defence evolution using a full spectrum of predator types.

Altogether, our results emphasize that immature birds, possibly due to their naivety, are less prone to detect cryptic prey compared to more experienced adults (Mappes et al. 2014). Overall, the effectiveness of crypsis in evading detection is evident, particularly in diverse environments where immature birds dominate, as in our study location in mature secondary forest habitat.

**Probability of being rejected upon identification:** Our results show that regardless of bird diet and species, predators often sight rejected *Heliconius* (i.e., assuming that predator and prey being closer than ∼10cm and there was no attack behaviour). This suggests that wild birds recognised the aposematic local patterns of unprofitable prey (e.g., Finkbeiner et al., 2014; Dell’aglio et al., 2016). Nevertheless, immature birds, presumably being more naïve, exhibited a less differentiated predation pattern, aligning with the findings of Mappes et al. (2014), where aposematism is more favourable during seasons with low numbers of naïve birds. In tropical habitats, naïve birds are present throughout year (Stouffer et al., 2013). However, this also creates an almost continuous selective pressure on aposematic signals throughout the year and may be linked to the evolution of diverse warning cues in the tropics.

In laboratory experiments, birds can learn phenotypic cues associated with escape ability and change their predation behaviour to avoid such evasive prey (Páez et al., 2021; Linke et al., 2022; Loeffler-Henry & Sherratt, 2024). Additionally, phenotypic cues commonly found in evasive neotropical butterflies, such as dorsal iridescence, forewing bands and hindwing tails, might advertise difficulty of capture (Janzen et al., 2009; Pinheiro & Freitas, 2014; Linke et al., 2022). While testing avoidance due to evasive capabilities has seldom been tested with wild predators (Guerra et al., 2024), our results hint at some avoidance by immature birds. Nonetheless, attack avoidance of evasive prey might be higher in species less adept at catching flying insects (such as Cardinalidae or Thraupidae) but due to our low sample size and unequal abundances of immatures and adults across habitats, this speculation remains to be corroborated.

**Targeting probability and capture avoidance:** Although there are slight variations in capture rates between immature and adult birds, these differences are relatively minor. This suggests that all three prey phenotypes were equally susceptible to be captured when attacked, regardless of the birds’ experience. This equal likelihood of capture could also be explained by the restricted space in our aviaries available for prey to escape capture. However, we found that adults were slightly better than immatures in catching the evasive prey *Spicauda*, as expected under the age-specific foraging proficiency concept (Wunderle, 1991).

Attack rates, on the other hand, diverge drastically between adult and immature birds. Adults preferred to attack *Spicauda* and refrained from attacking *Heliconius*, while immatures had similar attack rates towards all three prey phenotypes. Learning and memorizing aposematic cues of unprofitable prey are well-recognized by behavioural experiments in the laboratory (Gibson, 1974, 1980; Skelhorn & Rowe, 2006; Hansen et al., 2010; Páez et al., 2021; Zvereva & Kozlov, 2021; Linke et al., 2022), including tests for birds of different age classes (Langham, 2006; Veselý et al., 2017). Because the precise age determination of birds is impractical in the wild, immature individuals from our study may have encompassed a spectrum ranging from recently fledged to residents of up to two years. However, predatorial behaviours in wild birds were rarely assessed using age groups (Guerra et al., 2024), or were limited in sample size (Pinheiro & Campos, 2019). Thus, to our knowledge, age-related variation in prey attack and how it shapes the evolution and co-existence of multiple defences have only received little empirical support.

Contrary to the expectations of evasive ability importance in avoiding subjugation, *Spicauda* was not less likely to be caught by insectivorous birds than the other prey types. Nevertheless, the total instances of failed attacks are considerably higher for *Spicauda* compared to *Heliconius* or Euptychiines, pointing towards a bigger struggle for the birds to catch and subjugate *Spicauda* (Figure 3(c)). Food deprivation of birds prior to the experiment and the closed experimental space might have increased the mortality of *Spicauda* compared to natural conditions, as in nature unsuccessful attacks would be less likely to be followed by more attempts. However, high attack rates to evasive species might be explained by its high profitability, especially for agile predators. Evasive species with high flying speed typically have high wing loading (Le Roy et al., 2019), which can be driven by increased body mass. Thus, such evasive species might be still profitable to chase, though not for less agile or naïve predators.

**Probability to survive upon predator attack:** Mortality rates were relatively even across prey phenotypes and bird age groups. While log-likelihood tests detected some differences for immatures, they were completely absent for adults, hinting at *Heliconius* and *Spicauda* being less likely to be killed when attacked by immature birds. *Heliconius* are rejected after attacks due to chemical defences, while *Spicauda* is harder to catch and subjugate and less likely to be killed by less skilled predators. This is at least partly in agreement with our initial prediction, as species with high evasive capabilities or chemical defences are less likely to be killed by predators, at least for immature birds. Additionally, it points towards predation pressure being more equally distributed in more diverse habitats, which has been supported for aquatic habitats but less so for terrestrial, tropical ones (Chang & Todd, 2023).

Our findings demonstrate that multiple antipredator strategies—crypsis, aposematism, and evasive behaviour—can be equally effective for surviving in tropical forest ecosystems, albeit in different ways and at different stages of the predation process. The relatively similar mortality rate across prey types suggest that effective predation pressure is relatively even, allowing diverse prey strategies to diversify and coexist. This pattern reflects the complexity of tropical food webs, where the high diversity of both predators and prey likely promotes the persistence of multiple defensive adaptations. The role of predator age further underscores how variation in predator experience and behaviour shapes the frequency of prey defences. Altogether, these results highlight how predation contributes to the ecological and evolutionary processes that sustain the large diversity of antipredator defences found in the tropics.

## Acknowledgments

We thank the Gallusser-Ramirez family for offering their property and their invaluable help and advice during our experiments. We thank students who helped processing nearly 300 hours of video material: Tereza Látalová, Sara Turková, Eliška Macháňová and Barbora Bělovská. We extent our gratitude towards Leonardo Ré Jorge for advice in statistical analysis, Mario A. Marin for help in determining Euptychiines, Alena Sucháčková for comments on the manuscript, and Peruvian authorities at Servicio Nacional Forestal y de Fauna Silvestre – SERFOR (Ministry of Agriculture) and colleagues at the Departments of Ornithology and Entomology (Diana Silva and Gerardo Lamas) of the Natural History Museum, Universidad Nacional Mayor de San Marcos.

## Authors contributions

**DL** – study planning, field investigation, data curation and evaluation, video processing, statistical analysis, writing original draft, review and editing; **JHM** – field investigation, review and editing; **VNPNE** – field investigation, review and editing; **PG** – video processing, writing and editing; **CRG** – field investigation; **LSS** – study planning, supervision, review and editing; **ME** – study planning, statistical analysis, review and editing; **PMM** – study planning, field investigation, funding acquisition, review and editing.

## Funding

Funding was provided by the Junior GAČR grant (GJ20-18566Y), the PPLZ program of the Czech Academy of Sciences (fellowship grant L20096195) and GAJU n.014/2022/P. Logistic support in Peru was supported by 421 Fundación San Marcos, Letty Salinas was partially funded by Universidad Nacional Mayor de San Marcos RR 05557-R-22 (Project B22100321).

Birds used in this study were under the national permits D000088-2021-MIDAGRI-SERFOR-DGGSPFFS and D000047-2022-MIDAGRI-SERFOR-DGGSPFFS-DGSPFS. Butterflies were collected under the same national permits and RDG N.° 0488-2019-MINAGRI-SERFOR-DGGSPFFS and exported under the permits 003735-SERFOR and 003758-SERFOR.

## References

Arias, M., Mappes, J., Théry, M., & Llaurens, V. (2016). Inter-species variation in unpalatability does not explain polymorphism in a mimetic species. Evolutionary Ecology, 30(3), 419–433. 10.1007/s10682-015-9815-2

Bates, D., Mächler, M., Bolker, B., & Walker, S. (2015). Fitting Linear Mixed-Effects Models Using lme4. Journal of Statistical Software, 67, 1–48. 10.18637/jss.v067.i01

Burd, M. (1994). Butterfly wing colour patterns and flying heights in the seasonally wet forest of Barro Colorado Island, Panama. Journal of Tropical Ecology, 10(4), 601–610. 10.1017/S0266467400008270

Burnham, K. P., & Anderson, D. R. (2013). Model Selection and Inference: A Practical Information-Theoretic Approach. Springer Science & Business Media.

Chai, P. (1986). Field observations and feeding experiments on the responses of rufous-tailed jacamars (Galbula ruficauda) to free-flying butterflies in a tropical rainforest. Biological Journal of the Linnean Society, 29(3), 161–189. 10.1111/j.1095-8312.1986.tb01772.x

Chang, C., & Todd, P. A. (2023). Reduced predation pressure as a potential driver of prey diversity and abundance in complex habitats. Npj Biodiversity, 2(1), 1–5. 10.1038/s44185-022-00007-x

Chotard, A., Ledamoisel, J., Decamps, T., Herrel, A., Chaine, A. S., Llaurens, V., & Debat, V. (2022). Evidence of attack deflection suggests adaptive evolution of wing tails in butterflies. Proceedings of the Royal Society B: Biological Sciences, 289(1975), 20220562. 10.1098/rspb.2022.0562

Chouteau, M., Arias, M., & Joron, M. (2016). Warning signals are under positive frequency-dependent selection in nature. Proceedings of the National Academy of Sciences, 113(8), 2164–2169. 10.1073/pnas.1519216113

Chouteau, M., Dezeure, J., Sherratt, T. N., Llaurens, V., & Joron, M. (2019). Similar predator aversion for natural prey with diverse toxicity levels. Animal Behaviour, 153, 49–59. 10.1016/j.anbehav.2019.04.017

Dell’aglio, D. D., Stevens, M., & Jiggins, C. D. (2016). Avoidance of an aposematically coloured butterfly by wild birds in a tropical forest. Ecological Entomology, 41(5), 627–632. 10.1111/een.12335

Egg, A. B., Parker, T. A., O’Neill, J. P., Lane, D. F., Stotz, D. F., & Schulenberg, T. S. (2010). Birds of Peru: Revised and Updated Edition. Princeton University Press. https://muse.jhu.edu/pub/267/monograph/book/30246

Endler, J. A. (1991). Interactions between predator and prey. Behavioural Ecology, 169–196.

Feltmate, B. W., & Williams, D. D. (1989). A test of crypsis and predator avoidance in the stonefly Paragnetina media (Plecoptera: Perlidae). Animal Behaviour, 37, 992–999. 10.1016/0003-3472(89)90143-7

Finkbeiner, S. D., Briscoe, A. D., & Reed, R. D. (2014). Warning signals are seductive: Relative contributions of color and pattern to predator avoidance and mate attraction in Heliconius butterflies. Evolution, 68(12), 3410–3420. 10.1111/evo.12524

Gibson, D. O. (1974). Batesian mimicry without distastefulness? Nature, 250(5461), Article 5461. 10.1038/250077a0

Gibson, D. O. (1980). The role of escape in mimicry and polymorphism: I. The response of captive birds to artificial prey. Biological Journal of the Linnean Society, 14(2), 201–214. 10.1111/j.1095-8312.1980.tb00105.x

Guerra, T. J., Braga, R. F., Camarota, F., Neves, F. S., & Fernandes, G. W. (2024, March 5). *Avian predators avoid attacking fly-mimicking beetles: A field experiment on evasive mimicry using artificial prey* (world) [Research-article]. Https://Doi.Org/10.1086/730263. 10.1086/730263

Hansen, B. T., Holen, Ø. H., & Mappes, J. (2010). Predators use environmental cues to discriminate between prey. Behavioral Ecology and Sociobiology, 64(12), 1991–1997. 10.1007/s00265-010-1010-4

Janzen, D. H., Hallwachs, W., Blandin, P., Burns, J. M., Cadiou, J.-M., Chacon, I., Dapkey, T., Deans, A. R., Epstein, M. E., Espinoza, B., Franclemont, J. G., Haber, W. A., Hajibabaei, M., Hall, J. P. W., Hebert, P. D. N., Gauld, I. D., Harvey, D. J., Hausmann, A., Kitching, I. J., … Wilson, J. J. (2009). Integration of DNA barcoding into an ongoing inventory of complex tropical biodiversity. Molecular Ecology Resources, 9(s1), 1–26. 10.1111/j.1755-0998.2009.02628.x

Langham, G. M. (2006). Rufous-tailed jacamars and aposematic butterflies: Do older birds attack novel prey? Behavioral Ecology, 17(2), 285–290. 10.1093/beheco/arj027

Le Roy, C., Debat, V., & Llaurens, V. (2019). Adaptive evolution of butterfly wing shape: From morphology to behaviour. Biological Reviews, 94(4), 1261–1281. 10.1111/brv.12500

Linke, D., Elias, M., Klečková, I., Mappes, J., & Matos-Maraví, P. (2022). Shape of Evasive Prey Can Be an Important Cue That Triggers Learning in Avian Predators. Frontiers in Ecology and Evolution, 10. 10.3389/fevo.2022.910695

Linke, D., Hernandez Mejia, J., N. P. Eche Navarro, V., Salinas Sánchez, L., de Gusmão Ribeiro, P., Elias, M., & Matos-Maraví, P. (2024). Reduced palatability, fast flight, and tails: Decoding the defence arsenal of Eudaminae skipper butterflies in a Neotropical locality. Journal of Evolutionary Biology, voae091. 10.1093/jeb/voae091

Loeffler-Henry, K., & Sherratt, T. N. (2024). Selection for evasive mimicry imposed by an arthropod predator. Biology Letters, 20(1), 20230461. 10.1098/rsbl.2023.0461

Mallet, J., & Gilbert, L. (1995). Why are there so many mimicry rings? Correlations between habitat, behaviour and mimicry in Heliconius butterflies. Biological Journal of the Linnean Society, 55(2), 159–180. 10.1111/j.1095-8312.1995.tb01057.x

Mappes, J., Kokko, H., Ojala, K., & Lindström, L. (2014). Seasonal changes in predator community switch the direction of selection for prey defences. Nature Communications, 5(1), Article 1. 10.1038/ncomms6016

Mérot, C., Frérot, B., Leppik, E., & Joron, M. (2015). Beyond magic traits: Multimodal mating cues in Heliconius butterflies. Evolution, 69(11), 2891–2904. 10.1111/evo.12789

Molleman, F., Javoiš, J., Davis, R. B., Whitaker, M. R. L., Tammaru, T., Prinzing, A., Õunap, E., Wahlberg, N., Kodandaramaiah, U., Aduse-Poku, K., Kaasik, A., & Carey, J. R. (2020). Quantifying the effects of species traits on predation risk in nature: A comparative study of butterfly wing damage. Journal of Animal Ecology, 89(3), 716–729. 10.1111/1365-2656.13139

Navarro, V., Hernández Mejía, J., Linke, D., Salinas, L., & Matos-Maraví, P. (2023). Birds of Tarapoto (San Martín, Peru) (1640; p. 12). Field Guides, Field Museum. 10.13140/RG.2.2.14541.20964

Páez, E., Valkonen, J. K., Willmott, K. R., Matos-Maraví, P., Elias, M., & Mappes, J. (2021). Hard to catch: Experimental evidence supports evasive mimicry. Proceedings of the Royal Society B: Biological Sciences, 288(1946), 20203052. 10.1098/rspb.2020.3052

Pembury Smith, M. Q. R., & Ruxton, G. D. (2021). Size-dependent predation risk in cryptic prey. Journal of Ethology, 39(2), 191–198. 10.1007/s10164-021-00691-5

Pinheiro, C. E. G., & Campos, V. C. (2019). The responses of wild jacamars (Galbula ruficauda, Galbulidae) to aposematic, aposematic and cryptic, and cryptic butterflies in central Brazil. Ecological Entomology, 44(4), 441–450. 10.1111/een.12723

Pinheiro, C. E. G., & Cintra, R. (2017). Butterfly Predators in the Neotropics: Which Birds are Involved? Journal of the Lepidopterists’ Society. 10.18473/lepi.71i2.a5

Pinheiro, C. E. G., & Freitas, A. V. L. (2014). Some Possible Cases of Escape Mimicry in Neotropical Butterflies. Neotropical Entomology, 43(5), 393–398. 10.1007/s13744-014-0240-y

Ruxton, G. D., Allen, W. L., Sherratt, T. N., & Speed, M. P. (2018). Avoiding Attack: The Evolutionary Ecology of Crypsis, Aposematism, and Mimicry (2nd ed.). Oxford University Press. 10.1093/oso/9780199688678.001.0001

Ruxton, G. D., Speed, M., & Sherratt, T. N. (2004). Evasive mimicry: When (if ever) could mimicry based on difficulty of capture evolve? Proceedings. Biological Sciences, 271(1553), 2135–2142. 10.1098/rspb.2004.2816

Seymoure, B. M., Raymundo, A., McGraw, K. J., Owen McMillan, W., & Rutowski, R. L. (2018). Environment-dependent attack rates of cryptic and aposematic butterflies. Current Zoology, 64(5), 663–669. 10.1093/cz/zox062

Skelhorn, J., & Rowe, C. (2006). Prey palatability influences predator learning and memory. Animal Behaviour, 71(5), 1111–1118. 10.1016/j.anbehav.2005.08.011

Srygley, R. B., & Chai, P. (1990). Predation and the Elevation of Thoracic Temperature in Brightly Colored Neotropical Butterflies. The American Naturalist, 135(6), 766–787.

Stevens, M., & Merilaita, S. (2009). Animal camouflage: Current issues and new perspectives. Philosophical Transactions of the Royal Society B: Biological Sciences, 364(1516), 423–427. 10.1098/rstb.2008.0217

Stouffer, P., Johnson, E., & Bierregaard, R. (2013). Breeding Seasonality in Central Amazonian Rainforest Birds. The Auk, 130, 529–540. 10.1525/auk.2013.12179

Vallin, A., Jakobsson, S., Lind, J., & Wiklund, C. (2006). Crypsis versus intimidation—Anti-predation defence in three closely related butterflies. Behavioral Ecology and Sociobiology, 59(3), 455–459. 10.1007/s00265-005-0069-9

Veselý, P., Ernestová, B., Nedvěd, O., & Fuchs, R. (2017). Do predator energy demands or previous exposure influence protection by aposematic coloration of prey? Current Zoology, 63(3), 259–267. 10.1093/cz/zow057

Wickham, H. (2016). Ggplot2. Springer International Publishing. 10.1007/978-3-319-24277-4

Willmott, K. R., Robinson Willmott, J. C., Elias, M., & Jiggins, C. D. (2017). Maintaining mimicry diversity: Optimal warning colour patterns differ among microhabitats in Amazonian clearwing butterflies. Proceedings. Biological Sciences, 284(1855), 20170744. 10.1098/rspb.2017.0744

Wunderle, J., Joseph. (1991). Age-specific foraging proficiency in birds. Current Ornithology, 8, 273–324.

Zvereva, E. L., & Kozlov, M. V. (2021). Seasonal variations in bird selection pressure on prey colouration. Oecologia, 196(4), 1017–1026. 10.1007/s00442-021-04994-9

