## Supplement 1-8 for "Quantifying the success of prey crypsis, aposematism and evasiveness in avoiding predators’ attack"

**Supplement 1**: Coordinates of all nets used to trap low flying birds

*Table 1: Nets used to trap birds from the urban environment during the rainy season from October 2021 to February 2022.*

| **Name of the net** | **Longitude** | **Latitude** | **Altitude [masl]** | **Site description** |
| --- | --- | --- | --- | --- |
| casa | -76.35609576 | -6.478592891 | 380 | Forest edge |
| 1 casa | -76.35618622 | -6.478610737 | 381 | Forest edge |
| 2 casa | -76.35622227 | -6.478565422 | 383 | Forest edge |
| 3-4 casa | -76.35617694 | -6.478520325 | 382 | Forest edge |
| 5 casa | -76.35615893 | -6.478547504 | 382 | Forest edge |
| 6 casa | -76.35613420 | -6.479442893 | 370 | Forest edge |
| 7 casa | -76.35607095 | -6.479461150 | 368 | Forest edge |
| 8 casa | -76.35626054 | -6.479352119 | 373 | Forest edge |
| 9-10 casa | -76.35633302 | -6.479406187 | 374 | Forest edge |
| 1-2 shilcayo | -76.35547340 | -6.475772927 | 378 | Open grassland |
| 3-4-5 shilcayo | -76.35505776 | -6.475873517 | 360 | Open grassland |
| 1 suchiche | -76.35799049 | -6.483851238 | 351 | Relict forest |
| 2 suchiche | -76.35799063 | -6.483905499 | 351 | Relict forest |

*Table 2: Nets used to trap birds from the forest environment during the rainy season from October 2021 to February 2022 and during the dry season from June to September 2022.*

| **Name of the net** | **Longitude** | **Latitude** | **Altitude [masl]** | **Site description** |
| --- | --- | --- | --- | --- |
| 1 rio | -76.33184732 | -6.465299459 | 674 | Closed secondary forest |
| 2 rio | -76.33191080 | -6.465371643 | 673 | Closed secondary forest |
| piña chacra | -76.33513614 | -6.460986010 | 714 | Forest edge |
| 1-2 caseta | -76.33624607 | -6.463605785 | 724 | Low density secondary forest |
| 3 caseta | -76.33614585 | -6.463316648 | 723 | Closed secondary forest |
| 4 caseta | -76.33618128 | -6.463036199 | 724 | Closed secondary forest |
| 5 caseta | -76.33607416 | -6.463561018 | 721 | Closed secondary forest |
| 6-7 caseta | -76.33631840 | -6.463605595 | 725 | Low density secondary forest |
| tambo | -76.33347959 | -6.460230678 | 751 | Clearing of secondary forest |
| 1 chacra | -76.33379675 | -6.460501159 | 740 | Low density secondary forest |
| 2 chacra | -76.33394141 | -6.460500780 | 737 | Low density secondary forest |
| 3 chacra | -76.33413028 | -6.460120447 | 732 | Low density secondary forest |
| 4 chacra | -76.33401424 | -6.460690508 | 734 | Low density secondary forest |
| 5 chacra | -76.33409565 | -6.460708382 | 732 | Low density secondary forest |
| 6 chacra | -76.33423194 | -6.460961250 | 730 | Low density secondary forest |
| 7-8 chacra | -76.33383002 | -6.459397734 | 739 | Clearing of secondary forest |
| 9-10 chacra | -76.33376704 | -6.459515468 | 741 | Clearing of secondary forest |
| 11-12 chacra | -76.33361708 | -6.460944774 | 740 | Low density secondary forest |
| 13 chacra | -76.33370711 | -6.460799838 | 740 | Low density secondary forest |

**Supplement 2**: Photos of the aviaries in the urban and forested localities.

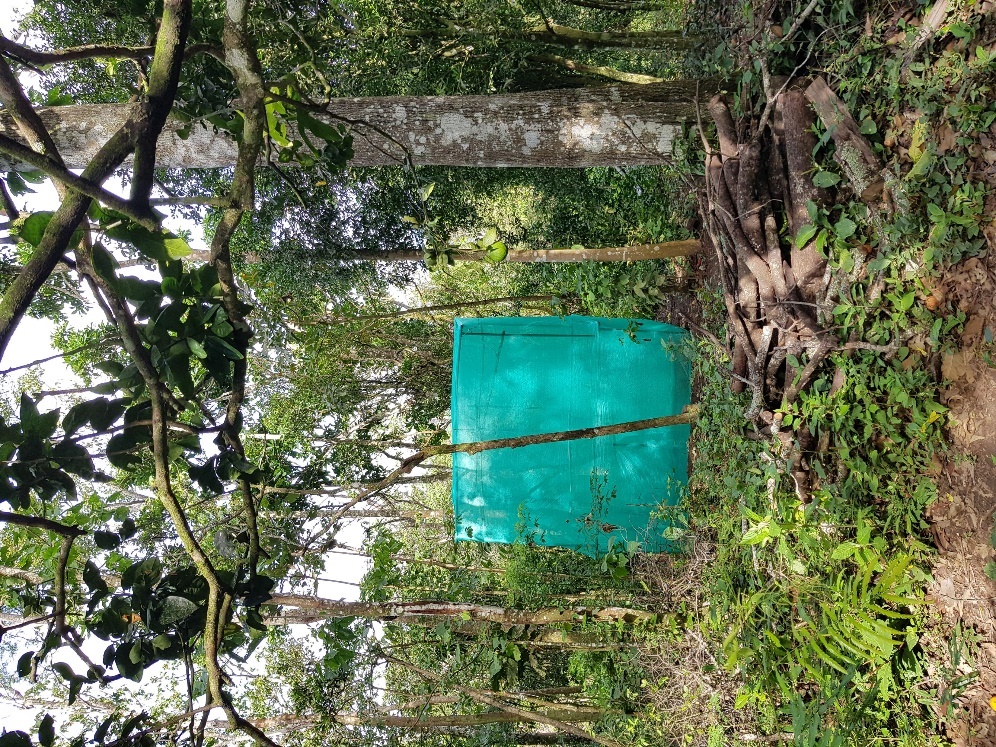

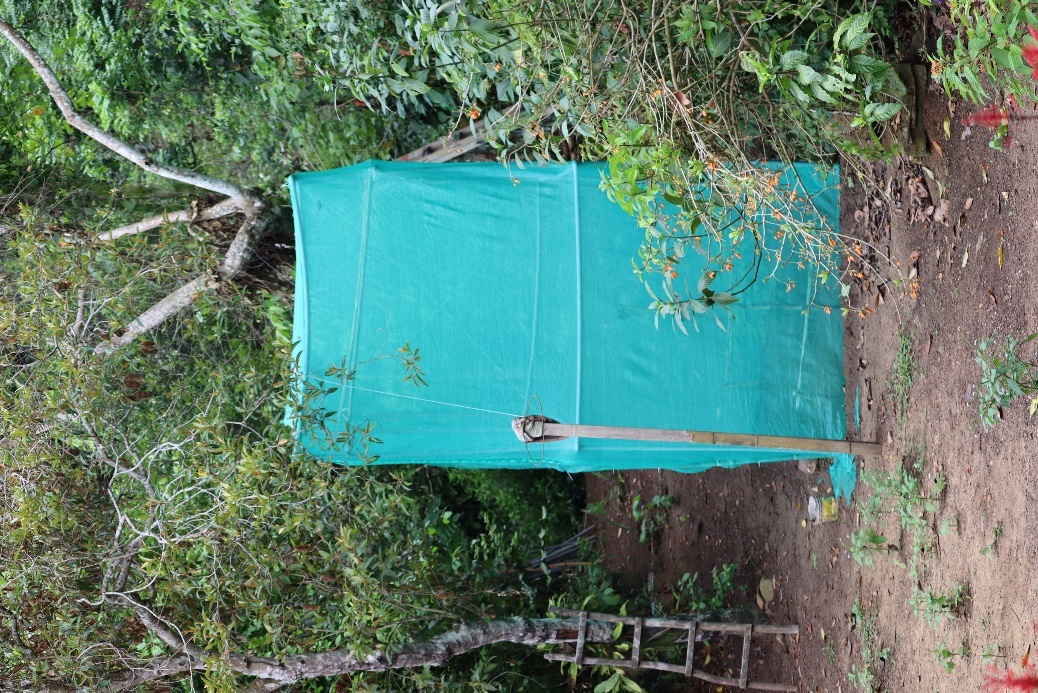
*Figure: Aviary at the urban (left) and forested (right) study sites. Outer dimensions approximately: height = 4m, depth = 2m and length = 4m. Tubing in the middle marked the 200 cm height used to separate the higher and lower stratum. Bamboo perch for bird to rest was installed approximately 50cm below the mesh ceiling.*

**Supplement 3**: Logistic regression models

The formulas “*sight rejection / attacking ~ butterfly_type + state + diet_reduced + (1|bird_ID)*” specified in generalized linear mixed models (GLMMs). The response variables, sight rejection or attacking, represented each a binary outcome (sight rejected or not; actively attacked or not) using the Family binomial (logit). Fixed effects included butterfly_type (type of butterfly; *Spicauda*, *Heliconius* or Euptychiina), state (bird age; juvenile or adult), and diet_reduced (diet category; insectivorous or other). The term (1|bird_ID) indicated a random effect for bird_ID (unique ID for each experimental bird), accounting for variability between individual birds.

***Ignoring behaviour***

```
Generalized linear mixed model fit by maximum likelihood (Laplace Approximation) ['glmerMod']
 Family: binomial  ( logit )
Formula: sight_rejection ~ butterfly_type + state + diet_reduced + (1|bird_ID)
   Data: data

     AIC      BIC   logLik deviance df.resid
   230.3    250.3   -109.2    218.3      202

Scaled residuals:
    Min      1Q  Median      3Q     Max
-44.057  -0.024   0.000   0.022  38.684

Random effects:
 Groups  Name        Variance Std.Dev.
 bird_ID (Intercept) 305.2    17.47
Number of obs: 208, groups:  bird_ID, 131

Fixed effects:
                         Estimate Std. Error z value Pr(>|z|)
(Intercept)               -7.9268     1.6213  -4.889 1.01e-06 ***
butterfly_typeHeliconius  14.7111     2.0900   7.039 1.94e-12 ***
butterfly_typeSpicauda    -0.1707     1.1547  -0.148    0.882
stateimmature              0.8333     1.1762   0.708    0.479
diet_reducedother          0.3889     1.1963   0.325    0.745
---
Signif. codes:  0 ‘***’ 0.001 ‘**’ 0.01 ‘*’ 0.05 ‘.’ 0.1 ‘ ’ 1

Correlation of Fixed Effects:
            (Intr) bttr_H bttr_S sttmmt
bttrfly_tyH -0.625
bttrfly_tyS -0.416  0.355
stateimmatr -0.382 -0.160  0.029
dit_rdcdthr -0.418 -0.119 -0.140  0.106
```

***Attacking behaviour***

```
Generalized linear mixed model fit by maximum likelihood (Laplace Approximation) ['glmerMod']
 Family: binomial  ( logit )
Formula: attacking ~ butterfly_type + state + diet_reduced + (1|bird_ID)
   Data: data

     AIC      BIC   logLik deviance df.resid
   438.6    462.4   -213.3    426.6      387

Scaled residuals:
    Min      1Q  Median      3Q     Max
-1.1320 -0.5327 -0.3691  0.6708  2.3807

Random effects:
 Groups  Name        Variance Std.Dev.
 bird_ID (Intercept) 0.8793   0.9377
Number of obs: 393, groups:  bird_ID, 131

Fixed effects:
                         Estimate Std. Error z value Pr(>|z|)
(Intercept)               -1.5276     0.4160  -3.672 0.000241 ***
butterfly_typeHeliconius  -0.2758     0.3333  -0.827 0.407999
butterfly_typeSpicauda     1.0421     0.3105   3.356 0.000791 ***
stateimmature             -0.1637     0.3271  -0.500 0.616832
diet_reducedother          0.1308     0.3379   0.387 0.698684
---
Signif. codes:  0 ‘***’ 0.001 ‘**’ 0.01 ‘*’ 0.05 ‘.’ 0.1 ‘ ’ 1

Correlation of Fixed Effects:
            (Intr) bttr_H bttr_S sttmmt
bttrfly_tyH -0.362
bttrfly_tyS -0.459  0.488
stateimmatr -0.494  0.005 -0.030
dit_rdcdthr -0.574 -0.002  0.005 -0.012
```

**Additional GLMM models with habitat, season and bird family as fixed effects**

The generalized linear mixed models (GLMMs) compared the likelihood of bird’s sight rejection or attacking butterflies, using each a binomial distribution with a logit link. It included as fixed effects the butterfly type (*Spicauda*, *Heliconius*, Euptychiina), bird state (juvenile, adult), diet (insectivorous, other), season (dry, wet), location (urban, forest), and bird family (see Supplement 5), while accounting for individual variation in birds as a random effect (1 | bird_ID).

*Sight rejection*

There was no influence of seasonality or habitat on sight rejection behaviour of birds, except for Vireonidae and Tyrannidae, which had lower likelihoods to sight reject butterflies compared to all other bird families. This interaction was significant for Vireonidae (z = -2.192, p = 0.028).

```
Generalized linear mixed model fit by maximum likelihood (Laplace Approximation) ['glmerMod']
 Family: binomial  ( logit )
Formula: sight_rejection ~ butterfly_type + state + diet + season + location + bird_family + (1 | bird_ID)
   Data: data

     AIC      BIC   logLik deviance df.resid
   263.6    327.0   -112.8    225.6      189

Scaled residuals:
     Min       1Q   Median       3Q      Max
-2.30868 -0.38146  0.09944  0.37237  2.07734

Random effects:
 Groups  Name        Variance Std.Dev.
 bird_ID (Intercept) 5.161    2.272
Number of obs: 208, groups:  bird_ID, 131

Fixed effects:
                           Estimate Std. Error z value Pr(>|z|)
(Intercept)                 0.19459    2.16904   0.090   0.9285
butterfly_typeHeliconius    3.17464    0.67485   4.704 2.55e-06 ***
butterfly_typeSpicauda      0.03910    0.68015   0.057   0.9542
stateimmature               0.71655    0.91515   0.783   0.4336
dietdother         	    -0.14868    0.94824  -0.157   0.8754
seasonwet                   0.07293    1.03887   0.070   0.9440
locationTarapoto            1.82380    1.80356   1.011   0.3119
bird_familyCotingidae      -0.25058    2.70196  -0.093   0.9261
bird_familyFurnariidae     -2.25679    1.86132  -1.212   0.2253
bird_familyIcteridae       -3.57406    3.62605  -0.986   0.3243
bird_familyPipromorphidae  -1.59743    1.70680  -0.936   0.3493
bird_familyThamnophilidae  -2.57102    2.06917  -1.243   0.2140
bird_familyThraupidae      -3.00807    2.35891  -1.275   0.2022
bird_familyTityridae       -1.60136    3.49774  -0.458   0.6471
bird_familyTroglodytidae  -13.90736  663.89990  -0.021   0.9833
bird_familyTurdidae        -1.94294    1.67428  -1.160   0.2459
bird_familyTyrannidae      -3.98502    2.38656  -1.670   0.0950 .
bird_familyVireonidae      -5.38696    2.45800  -2.192   0.0284 *
---
Signif. codes:  0 ‘***’ 0.001 ‘**’ 0.01 ‘*’ 0.05 ‘.’ 0.1 ‘ ’ 1
```

*Attacking behaviour*

There was no influence of season but almost significant influence of location, with the Urban area (Tarapoto) showing higher attack rates compared to the forested area. This might be due to a higher number of adult birds in the urban area. Tyrannidae and Vireonidae showed higher probabilities to attack butterflies compared to Cardinalidae.

```
Generalized linear mixed model fit by maximum likelihood (Laplace Approximation) ['glmerMod']
 Family: binomial  ( logit )
Formula: attacking ~ butterfly_type + state + diet + season + location + bird_family + (1 | bird_ID)
   Data: data

     AIC      BIC   logLik deviance df.resid
   453.2    528.7   -207.6    415.2      374

Scaled residuals:
    Min      1Q  Median      3Q     Max
-1.7123 -0.5355 -0.3904  0.6533  2.9532

Random effects:
 Groups  Name        Variance Std.Dev.
 bird_ID (Intercept) 0.6142   0.7837
Number of obs: 393, groups:  bird_ID, 131

Fixed effects:
                          Estimate Std. Error z value Pr(>|z|)
(Intercept)                -2.4361     1.0972  -2.220 0.026397 *
butterfly_typeHeliconius   -0.2943     0.3338  -0.881 0.378063
butterfly_typeSpicauda      1.0185     0.3083   3.303 0.000956 ***
stateimmature              -0.1153     0.4377  -0.263 0.792283
dietother                   0.1281     0.4673   0.274 0.784025
seasonwet                   0.4869     0.5168   0.942 0.346175
locationTarapoto           -1.3921     0.7481  -1.861 0.062758 .
bird_familyCotingidae      -0.2359     1.5150  -0.156 0.876241
bird_familyFurnariidae      1.3629     0.9321   1.462 0.143689
bird_familyIcteridae        1.0549     1.6821   0.627 0.530561
bird_familyPipromorphidae   0.3942     0.8648   0.456 0.648479
bird_familyThamnophilidae   0.8205     1.0310   0.796 0.426125
bird_familyThraupidae       1.6570     1.1173   1.483 0.138058
bird_familyTityridae        0.8594     1.8084   0.475 0.634645
bird_familyTroglodytidae    1.4397     1.8186   0.792 0.428567
bird_familyTurdidae         0.6864     0.8431   0.814 0.415565
bird_familyTyrannidae       2.0372     1.0895   1.870 0.061517 .
bird_familyVireonidae       2.3088     1.0935   2.111 0.034744 *
---
Signif. codes:  0 ‘***’ 0.001 ‘**’ 0.01 ‘*’ 0.05 ‘.’ 0.1 ‘ ’ 1
```

**Supplement 4:** Results of log-likelihood tests for habitat and season.

*Table 1: Likelihood-based model tests of scenarios depicting the effectiveness of antipredator defences by three butterfly prey along the predation sequence. Akaike weights for the total number of birds, season (dry vs. wet) and habitat (urban vs. forest) are shown. Best models are marked in* ***bold and underlined****, while competing models (ΔAICc < 2) are only* ***bold****.* *Values in brackets represent ΔAICc values compared to the best performing model.*

| **Stage** | **Model** | **All different** | **All equal** | ***Heliconius* different** | ***Spicauda* different** | **Euptychiina different** |
| --- | --- | --- | --- | --- | --- | --- |
| (1) Remaining undetected | Total | **0.47** (0.18) | 0.00 (15.45) | 0.02 (6.98) | 0.00 (16.58) | **0.51** (0.00) |
|  | Urban | **0.15** (1.25) | **0.22** (0.51) | 0.08 (2.58) | **0.28** (0.02) | **0.28** (0.00) |
|  | Forest | **0.72** (0.00) | 0.00 (16.55) | 0.13 (3.47) | 0.00 (18.56) | 0.15 (3.10) |
|  | Dry season | **0.53** (0.00) | 0.00 (12.65) | **0.41** (0.50) | 0.00 (14.44) | 0.06 (4.42) |
|  | Wet season | **0.24** (1.94) | 0.04 (5.70) | 0.02 (6.50) | 0.05 (4.93) | **0.64** (0.00) |
| (2) Rejection upon identification | Total | **0.50** (0.00) | 0.00 (26.34) | **0.50** (0.02) | 0.00 (25.15) | 0.00 (13.10) |
|  | Urban | 0.23 (2.04) | 0.03 (5.97) | **0.65** (0.00) | 0.05 (5.09) | 0.03 (6.22) |
|  | Forest | **0.62** (0.00) | 0.00 (19.54) | **0.37** (1.04) | 0.00 (20.32) | 0.01 (7.53) |
|  | Dry season | **0.40** (0.76) | 0.00 (12.91) | **0.58** (0.00) | 0.00 (13.39) | 0.02 (6.49) |
|  | Wet season | **0.34** (1.21) | 0.00 (11.68) | **0.63** (0.00) | 0.00 (12.06) | 0.02 (6.64) |
| (3.1) Targeting probability | Total | **0.25** (1.77) | 0.04 (5.73) | 0.07 (4.29) | **0.62** (0.00) | 0.02 (6.70) |
|  | Urban | **0.50** (0.00) | 0.00 (9.32) | 0.17 (2.16) | **0.32** (0.91) | 0.00 (11.03) |
|  | Forest | 0.10 (2.45) | **0.33** (0.00) | **0.13** (1.83) | **0.26** (0.46) | **0.17** (1.34) |
|  | Dry season | 0.08 (3.25) | **0.39** (0.00) | 0.14 (2.02) | **0.18** (1.57) | **0.21** (1.26) |
|  | Wet season | **0.36** (0.54) | 0.01 (6.99) | 0.14 (2.49) | **0.48** (0.00) | 0.01 (8.64) |
| (3.2) Capture avoidance | Total | 0.09 (2.89) | **0.37** (0.00) | **0.16** (1.66) | **0.14** (1.88) | **0.24** (0.86) |
|  | Urban | 0.06 (3.77) | **0.42** (0.00) | **0.18** (1.62) | **0.18** (1.67) | **0.15** (2.00) |
|  | Forest | 0.08 (3.23) | **0.39** (0.00) | **0.17** (1.62) | 0.14 (2.06) | **0.22** (1.17) |
|  | Dry season | 0.09 (2.80) | **0.35** (0.00) | 0.13 (2.05) | **0.15** (1.64) | **0.28** (0.45) |
|  | Wet season | 0.06 (4.13) | **0.45** (0.00) | 0.16 (2.03) | 0.16 (2.11) | **0.17** (1.98) |
| (4) Surviving attack | Total | 0.06 (3.79) | **0.43** (0.00) | **0.16** (1.99) | **0.16** (1.97) | **0.19** (1.67) |
|  | Urban | 0.06 (3.96) | **0.44** (0.00) | **0.21** (1.48) | 0.14 (2.26) | 0.14 (2.29) |
|  | Forest | 0.07 (3.32) | **0.39** (0.00) | 0.14 (2.11) | **0.19** (1.50) | **0.21** (1.27) |
|  | Dry season | 0.06 (4.20) | **0.45** (0.00) | **0.17** (1.97) | 0.15 (2.24) | **0.17** (1.93) |
|  | Wet season | 0.06 (3.93) | **0.43** (0.00) | **0.16** (1.97) | **0.18** (1.74) | **0.16** (2.00) |

**Supplement 5**: List of bird species used for valid experiments depending on habitat, season and age.

*Table 1: Breakdown of bird species used in successful experiments according to bird family and experimental season / habitat. Number in brackets represents the number of adult and immature individuals (Syntax = Adult-Immature). Diets classes were condensed for GLMM models to insectivorous and other (comprising all other diet classes).*

|  | **Urban-wet** | **Forest-wet** | **Forest-dry** | **Diet** |
| --- | --- | --- | --- | --- |
| **Bucconidae** 1 (1-0) | | | | |
| *Monasa morphoeus* |  |  | 1 (1-0) | insectivorous |
| **Cardinalidae** 8 (1-7) | | | | |
| *Chlorothraupis frenata* |  |  | 1 (0-1) | frugivore |
| *Habia rubica* |  | 5 (0-5) | 1 (1-0) | frugivore/invertebrate |
| *Piranga olivacea* | 1 (0-1) |  |  | frugivore/invertebrate |
| **Cotingidae** 2 (0-2) | | | | |
| *Lipaugus vociferans* |  | 2 (0-2) |  | frugivore/invertebrate |
| **Furnariidae** 20 (2-18) | | | | |
| *Dendrocincla fuliginosa* |  | 1 (0-1) | 3 (0-3) | insects/omnivore |
| *Dendroma erythorptera* |  | 1 (0-1) |  | omnivore |
| *Furnarius leucopus* | 1 (1-0) |  |  | insectivorous |
| *Glyphorynchus spirurus* |  |  | 3 (0-3) | insectivorous |
| *Sittasomus griseicapillus* |  |  | 1 (0-1) | frugivore/invertebrate |
| *Xenops minutus* |  |  | 5 (0-5) | insectivorous |
| *Xiphorhynchus elegans* |  | 2 (1-1) | 3 (0-3) | insects/omnivore |
| **Icteridae** 2 (2-0) | | | | |
| *Cacicus cela* | 2 (2-0) |  |  | insects/omnivore |
| **Pipromorphidae** 50 (8-42) | | | | |
| *Leptopogon amaurocephalus* |  | 9 (4-5) | 11 (2-9) | insectivorous |
| *Mionectes oleagineus* |  | 5 (2-3) | 24 (0-24) | frugivore/invertebrate |
| *Mionectes olivaceus* |  |  | 1 (0-1) | frugivore/invertebrate |
| **Thamnophilidae** 24 (12-12) | | | | |
| *Hypocnemis peruviana* |  |  | 1 (0-1) | insectivorous |
| *Myrmelastes leucostigma* |  |  | 1 (1-0) | insectivorous |
| *Myrmoborus myotherinus* |  | 2 (0-2) | 1 (1-0) | insectivorous |
| *Myrmotherula axillaris* |  |  | 6 (2-4) | insectivorous |
| *Pithys albifrons* |  |  | 2 (0-2) | insectivorous |
| *Sciaphylax hemimelaena* |  | 2 (2-0) | 2 (2-0) | frugivore/invertebrate |
| *Thamnophilus schistaceus* |  | 1 (1-0) | 2 (0-2) | insectivorous |
| *Thamnophilus unicolor* |  | 1 (1-0) |  | frugivore |
| *Willisornis poecilinotus* |  | 1 (1-0) | 2 (1-1) | insectivorous |
| **Thraupidae** 27 (25-2) | | | | |
| *Saltator maximus* |  | 1 (1-0) | 2 (1-1) | frugivore/invertebrate |
| *Thraupis episcopus* | 18 (17-1) |  |  | frugivore/invertebrate |
| *Thraupis palmarum* | 6 (6-0) |  |  | frugivore/invertebrate |
| **Tityridae** 2 (2-0) | | | | |
| *Myiobius barbatus* |  | 1 (1-0) |  | insectivorous |
| *Pachyramphus polychopterus* |  | 1 (1-0) |  | insectivorous |
| **Troglodytidae** 1 (0-1) | | | | |
| *Microcerculus marginatus* |  |  | 1 (0-1) | insectivorous |
| **Turdidae** 32 (0-32) | | | | |
| *Catharus ustulatus* | 3 (0-3) | 29 (0-29) |  | frugivore/invertebrate |
| **Tyrannidae** 38 (21-17) | | | | |
| *Attila spadiceus* |  | 3 (3-0) |  | omnivore |
| *Elaenia parvirostris* | 1 (1-0) |  |  | insects/omnivore |
| *Myiodynastes maculatus* |  |  | 2 (0-2) | insects/omnivore |
| *Myiozetetes similis* | 3 (3-0) |  |  | frugivore/invertebrate |
| *Pitangus sulphuratus* | 23 (12-11) |  |  | omnivore |
| *Rhynchocyclus olivaceus* |  |  | 5 (1-4) | insectivorous |
| *Tyrannus melancholicus* | 1 (1-0) |  |  | insectivorous |
| **Vireonidae** 9 (3-6) | | | | |
| *Vireo flavoviridis* | 3 (0-3) | 1 (0-1) |  | frugivore/invertebrate |
| *Vireo olivaceus* | 3 (1-2) | 2 (2-0) |  | frugivore/invertebrate |

**Supplement 6:** Overview of bird-butterfly-interactions along the predation sequence (Encounter, sight-rejection, attacked, caught, killed) depending on bird diet (Insectivorous vs. Other) and bird family

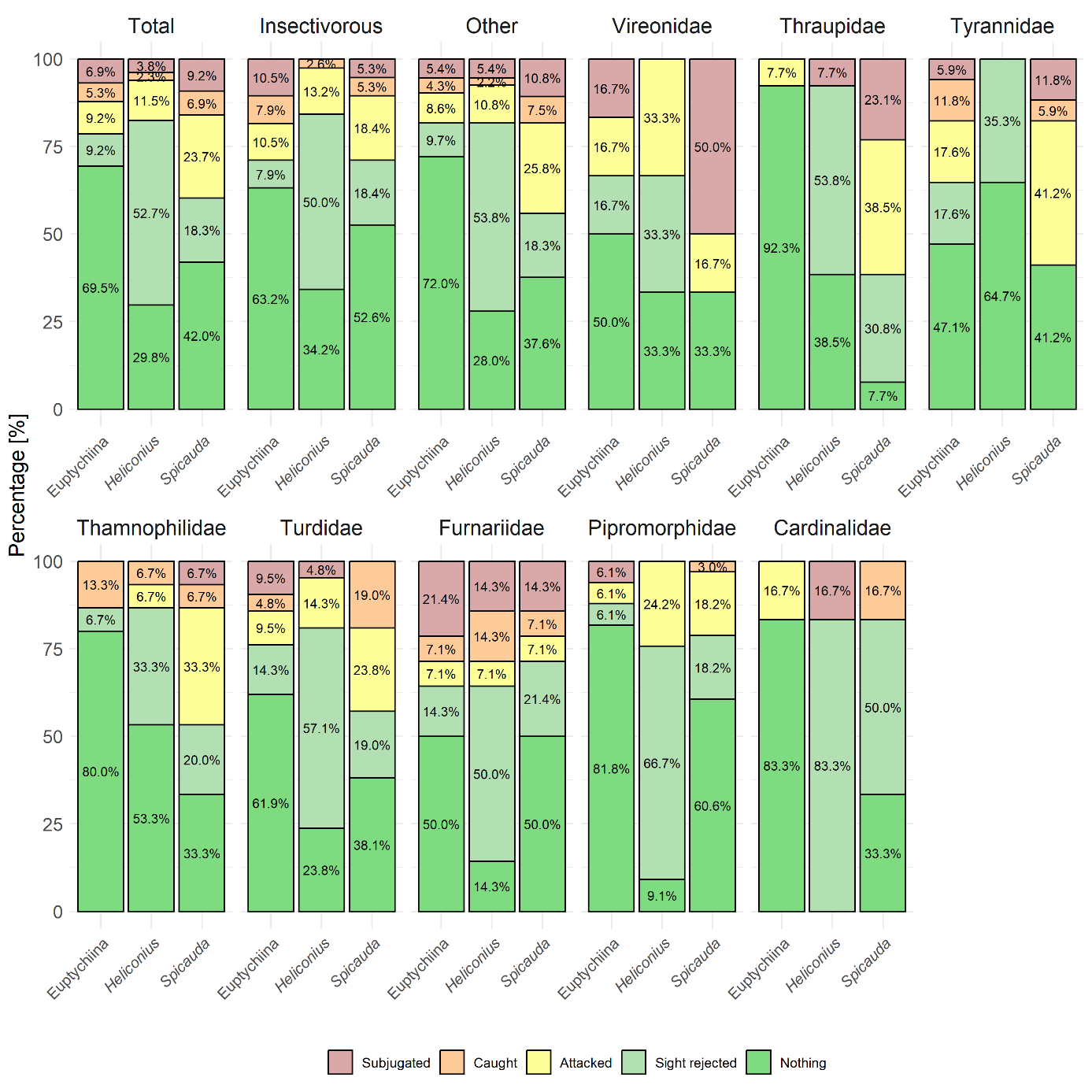

*Figure: Overview of the highest predator-prey-interaction (%) observed between prey phenotypes depending on the diet and bird family. Results for bird families are sorted from the highest attacking behaviour towards Spicauda to lowest. Only including families with 5 or more birds and birds which interacted with at least one prey phenotype, total and diet (insectivorous and other) includes bird families with less than 5 tested individuals.*

**Supplement 7:** Outcomes of predation experiments depending on habitat (Forest vs. Urban) season (Dry vs. Wet)
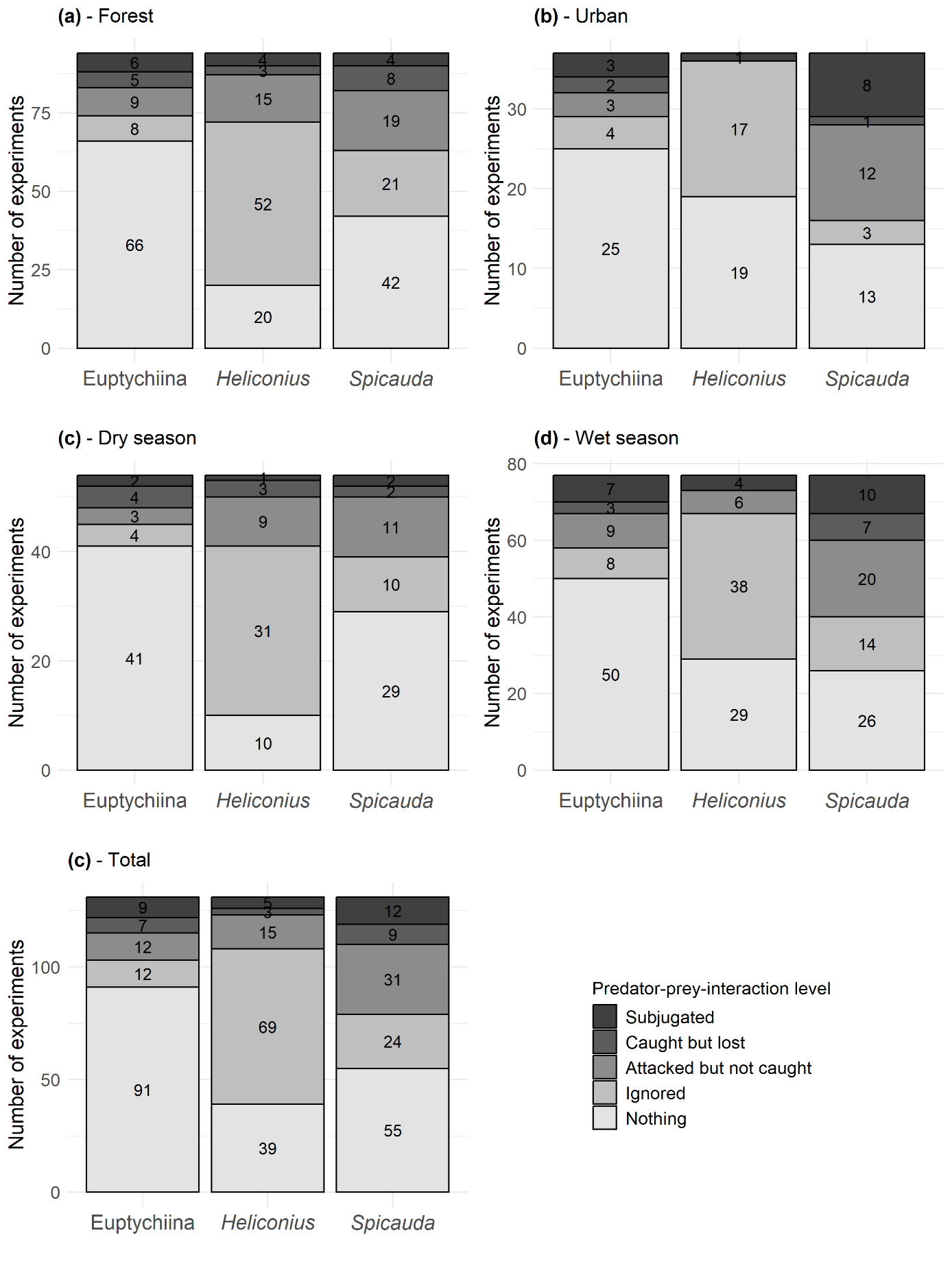

*Figure: Comparison of predation outcomes (nothing, sight rejected after detection, attacked, caught or subjugated) depending on the butterfly type (Euptychiines, Heliconius and Spicauda), season (dry and wet) and habitat (urban or forest).* ***(a)****: distribution for birds caught in the forested environment;* ***(b)****: distribution for all birds from the urban environment;* ***(c)****: distribution for all birds caught during the dry season;* ***(d)****: distribution for all birds caught during the wet season and* ***(e)****: distribution of the total dataset for comparison. Only the highest observed predation stage per bird-butterfly pair was used. Numbers inside bars indicate sample size.*

**Supplement 8:** Stratum selection in the aviaries (high > 200 cm, low < 200 cm) by bird families and butterfly type.
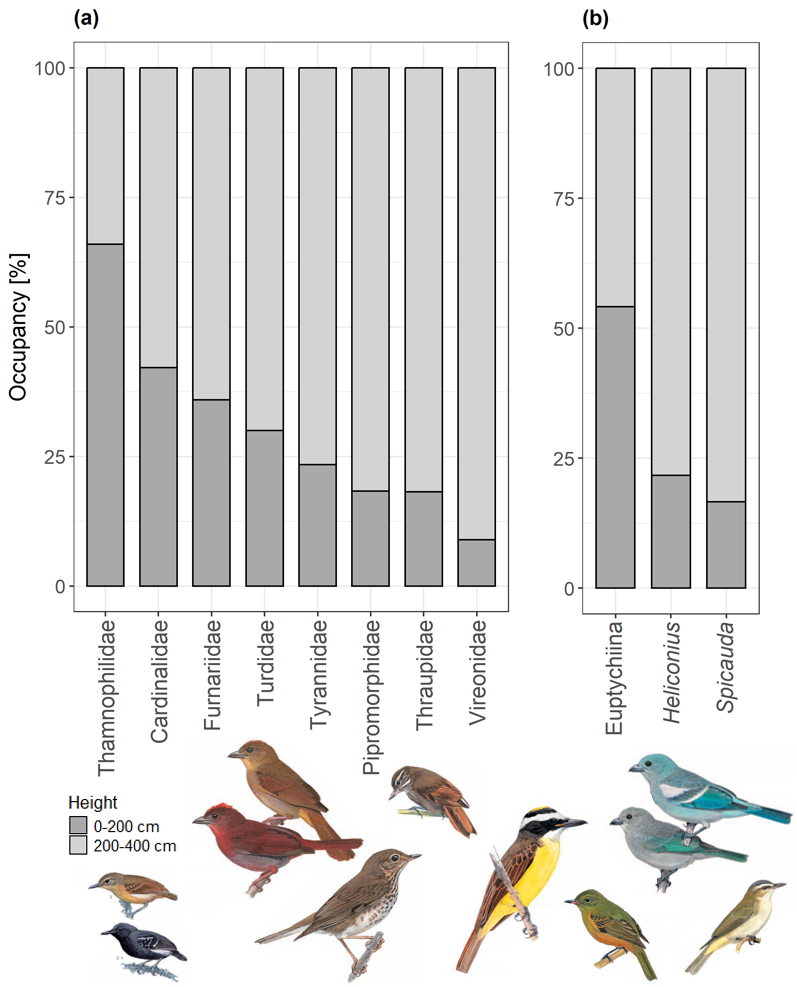

*Figure: Stratum preference of birds and butterflies.* ***(a)*** *Percentage of time spent (occupancy) by birds in the bottom, 0–200cm and the top, 200–400cm, of the aviary during the experiment (60 minutes). In the graph, only bird families with at least 5 valid experiments are presented. Drawing of birds are taken from Egg et al. (2010) and in order of panel A; from left to right: Myrmotherula axillaris, Habia rubica, Catharus ustulatus, Xenops minutus, Pitangus sulphuratus, Mionectes oleaginous, Thraupis episcopus and Vireo olivaceus. Bird pictures are scaled approximately.* ***(b)*** *Percentage of time spent by the three prey phenotypes (Euptychiines, Heliconius and Spicauda)* *at the two different heights in the aviary.*
